# Viscoelastic Deformability Cytometry: Ultra-high Throughput Platform for Mechanical Phenotyping of Cells in Liquid and Solid Biopsies

**DOI:** 10.1101/2023.11.17.567557

**Authors:** Mohammad Asghari, Sarah Duclos Ivetich, Mahmut Kamil Aslan, Morteza Aramesh, Melkonyan Oleksandr, Yingchao Meng, Rong Xu, Monica Colombo, Tobias Weiss, Stefan Balabanov, Stavros Stavrakis, Andew J. deMello

## Abstract

Due to the differences in mechanical properties between cancer cells and their benign counterparts, the mechanical characteristics of cells have emerged as a significant feature in diagnosing cancer. Consequently, measuring cell deformability as a means of mechanical phenotyping is promising for both the detection and classification of the disease. However, performing high-throughput single-cell deformability measurements on liquid or solid tissue biopsies remains a significant challenge within a clinical setting. Herein, we introduce an ultra-high throughput viscoelastic-based microfluidic platform to measure the cell mechanical properties at rates of up to 100,000 cells/s. Thanks to the viscoelastic property of the fluid, cells are focused and deformed within the same platform obviating the need for any sheath fluid. We used the presented platform for cell phenotyping in both liquid and solid tumor biopsies, such as identification of malignant lymphocytes in peripheral blood samples and glioblastoma-type cell classification from solid tumor samples. The presented platform has the potential to open new opportunities in the assessment of cancer, enable precise mechanical profiling of rare cells, and facilitate sensitive diagnostic applications.

## Introduction

Over the past decade, invasive techniques for diagnosing and monitoring cancers are slowly being replaced by non-invasive methods such as liquid biopsies. Liquid biopsies are minimally invasive and provide a methodology for obtaining tumor-derived information from body fluids. Much attention has focused on the molecular characterization of cells, intending to unravel biomarkers and detect various types of diseases. Isolation of cells can be performed by a variety of methods including capturing cells using specific biomarkers expressed on the cell surface. However, cancer cells can exhibit significant heterogeneity leading to variations in marker expression between different cells or at different stages of cancer progression. This heterogeneity can make it difficult to rely solely on specific markers for detection.

Cells are viscoelastic materials and undergo changes in their mechanical phenotype due to applied stress, caused during diverse physiological and disease processes. Thus, measuring cell mechanical properties such as cell deformability can be used as a biophysical marker for diseases inducing changes in subcellular constituents. For example, the deformability of single cells has been proposed as a label-free biomarker to enhance both disease diagnosis and clinical decision-making.^1, 2^ Altered cell deformability is especially relevant for cancer, where cancerous cells are typically more deformable than non-cancerous cells and can therefore be differentiated from healthy cells by their mechanical properties. ^3-7^ Whilst the origins of cell deformability in cancer are still poorly understood, its measurement has undoubted utility in cancer classification and diagnosis^2^ as well as developing new diagnosis methods.

To investigate structural alteration at the single-cell level, various techniques have been introduced, such as atomic force microscopy (AFM),^8^ gradual micropore filtering,^9^ micropipette aspiration,^10^ optical stretching,^11^ and magnetic tweezers.^12^ Even though these approaches provide accurate deformability values, they have very low analytical throughput (only a few cells per second) and require tedious operation procedures for both sample preparation and data acquisition, thus limiting their use only to proof-of-concept demonstrations. The widespread adoption of cell deformability measurements in clinical settings can be accelerated by the availability of high-throughput methods able to efficiently probe clinical samples containing large numbers of cells.

In this respect, microfluidic systems are particularly intriguing since recently they have emerged as an attractive alternative for studying cell mechanobiology and phenotyping.^13^ These approaches enable high throughput, offer integration with other detection methods, require minimal sample/reagent volumes, and provide good biocompatibility. Microfluidic systems for cell deformation analysis typically leverage either physical structures or fluid flow to induce cellular deformation. So far, three types of microfluidics-based approaches for measuring cell mechanical phenotypes have been reported as constriction-based deformability cytometry (cDC),^14-17^ shear flow deformability cytometry (sDC),^18-20^ and extensional flow deformability cytometry (xDC)^21-23^. Constriction-based deformability cytometry (cDC) relies on driving cells through a constriction smaller than the average diameter of the cells. By measuring the time cells need to pass through the constriction, the deformability of cells is directly deduced from their passage time. Nonetheless, the main disadvantage of this method is that the cells come into contact with the walls of the channel, which may result in blockage if a greater number of cells enter the channel.^17^ In contrast, both shear flow deformability cytometry (sDC) and extensional flow deformability cytometry (xDC) operate in a contactless manner by employing microfluidic channels with cross-sectional dimensions that exceed the average diameter of the cells. More specifically they utilize hydrodynamic forces to induce cell deformation and calculate cell deformability from image-based evaluation of cell shape.

A major difference between these two approaches is the use of different geometries to probe deformation thus having different timescales. In xDC, cells are stretched by placing them at a junction where two counter propagating liquids flow encounter.^21^ The cells are delivered to the junction at several meters per second and are fully decelerated and deformed within a few microseconds. This enables mechanical phenotyping of cells at a throughput up to 1000 events/s. Despite its effectiveness, the clinical application of xDC has been limited due to the complexity of fluid control, as it requires the balancing of two counter-propagating flows, and the necessity to conduct image analysis post-experiment.

In sDC, the cells are driven and focused into a one-dimensional array with the use of a sheath flow to a confining microfluidic channel with a funnel-like constriction geometry^18^. Shear forces and pressure gradients deform the cells into a bullet-like shape within the constriction region without contacting the channel walls. The sheath flow typically requires higher flow rates than the sample flow, a typical ratio being 3:1 to focus the cells centrally and apply enough shear stresses. Using the sDC approach, real-time throughputs of 100 cells/s are achievable. However, and as with conventional flow cytometers, this microfluidic system consumes large sample and reagent volumes due to a reliance on sheath flows for sample focusing. This means that parallelization is difficult to be implemented for a typical large-scale single-cell diagnosis application. Although both aforementioned approaches are non-invasive and allow for label-free analysis of mechanical properties, the clinical utility of single-cell deformability measurements is far from widespread either by a lack of real-time capability or low-throughput operation.

Recently, non-Newtonian viscoelastic fluids have attracted significant attention due to their diverse cell manipulation abilities such as focusing^24-26^ and deformation^27-29^. These fluids are prepared by simply dissolving biological or synthetic polymeric polymers inside the Newtonian fluids. The sDC microfluidic system, as with conventional flow cytometers, consumes large sample and reagent volumes due to a reliance on sheath flows for sample focusing. This means that parallelization is difficult and thus large-scale single cell diagnosis unfeasible. Viscoelastic fluids align the particles into a single file in the center of the channel, alleviating the need for using sheath flow for focusing and achieving single-plane focusing of cells in parallel microchannels prior to deformation.^30, 31^ Another advantage is that elastic forces and shear stresses on the flowing particles, can be used for single-file focusing and deformation of the cells in a single microfluidic device.^26^ More specifically viscoelastic fluids produce a symmetric shear stress distribution around the cells that can manipulate and deform the cells. Additionally, the elastic properties of the viscoelastic medium allow the same cell deformation to be achieved at a comparable linear velocity to a Newtonian carrier fluid.

These insights motivated us to create a real-time microfluidic platform that utilizes viscoelastic fluids to focus and deform cells within parallel microchannels without the need for sheath fluids. We term the method viscoelastic deformability cytometry (VDC), since we use viscoelastic microfluidics to both focus and deform cells and high-speed imaging to extract information related to cell deformation properties. High-resolution brightfield images of large numbers of single cells are obtained using a simple optical microscope and a CMOS camera sensor, with real-time image-processing capability used to quantify deformability and allow for label-free disease diagnosis. We first validated our VDC platform by measuring differences in the mechanical phenotype between different types of blood cells and breast cancer cell lines, BT474 and MDA-MB468 cells. Next we demonstrate the sensitivity of VDC in detecting changes in Jurkat cell deformability resulting from treatments with cytoskeletal-perturbing drugs. Furthermore, we utilized the VDC platform to successfully differentiate subpopulations of solid tumor derived cancer cells, obviating the need for molecular labeling. More specifically, we investigated the mechanical differences between two types of patient-derived glioma cells: glioma-initiating cells (GIC) that have stem-like properties and are resistant to chemo- and radiotherapy and more differentiated long-term serum-cultured cell lines (LTC). Additionally, we showcased the ability of our viscoelastic deformability to enable high throughput and sensitive diagnosis of chronic lymphomic leukemia (CLL). Thanks to the device’s simplicity and ultra-high throughput, the system has the potential to be widely used in clinical and research settings, particularly for diagnostic purposes.

## Results and Discussion

**Figure 1a** represents a schematic view of the viscoelastic deformability platform. The microfluidic device is simple in construction and has two main components: the first is responsible for cell focusing and the second one for inducing cell deformation. In the first component, cells flowing along a microfluidic channel are focused laterally and vertically due to elastic forces originating from a viscoelastic fluid. In the second component, flow through constriction channels induces high stresses on cells that subsequently causing deformation. In the focusing part of the system, each microchannel has a square cross-section (50 x 50 µm^2^) and a focusing length of 3 cm. In the deformation part, each narrow channel has a 15 x 15 µm^2^ cross-section and a length of 300 µm. Within the constricted channels, suspended cells are subject to both shear and normal stresses since the flow velocity is higher in the center of the channel when compared to regions closer to the channel walls. Such stresses cause cells to adopt “bullet-like” cell shapes, as predicted by computational fluid dynamics (CFD) simulations (**Figure 1b)** and shown in images of the deformation region (**Figure 1c).** To capture blur-free images of cells, exposure times were set to 2 µs with a recording rate of 10,000 frames per second. Single-cell images at the end of the deformation region, are used to measure the cell size (*A*) and deformability using the formula: 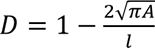, where *l* is the cell perimeter. Accordingly, scatter plot of cell deformability versus size can be constructed as depicted in **Figure 1d**. Since no sheath flow is required in VDC, the method can be readily parallelized by arranging multiple deformation microchannels in parallel, enabling simultaneous observation of cells in the microchannels (**Figure 1e**). Advantageously, the cells in multiple deformation microchannels are imaged by the same imaging device. In this manner, cells can be imaged and analyzed at extremely high throughput, e.g., at up to 100’000 cells/second, a value two orders of magnitude higher than the current state-of-the-art.^32^

**Figure 1.**
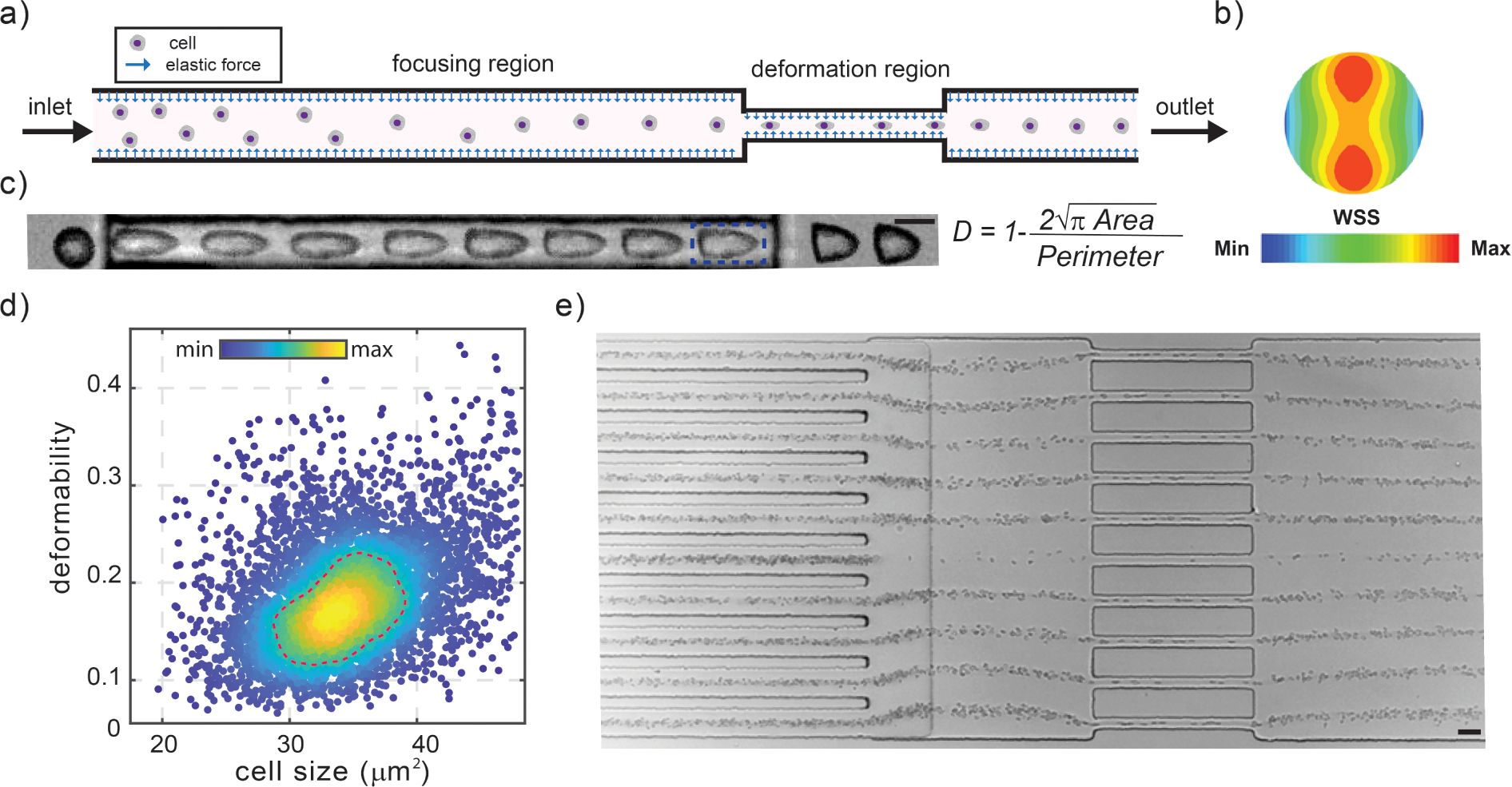
Image-based Viscoelastic Deformability Cytometry (VDC). (a) The operational principle of the VDC platform. Randomly distributed cells entering the focusing region are centrally focused due to elastic forces. Next, they enter a constriction region where they deform and finally relax before exiting at the outlet. b) Shear stress distribution around the cell inside the constriction channel. Surface color indicates magnitude. c) Magnified image of the deformation region showing the progression of cell deformability along the channel length (scale bar: 10µm). Image of the cell deformation is acquired in the region marked by the blue dotted line. d) A representative scatter plot of deformability vs cell size extracted from measurements of red blood cells. Color indicates a linear density scale; dotted line, 50%-density contour. e) Focusing and deformation of cells in the VDC microfluidic device consisting an array of parallel channels (scale bar: 30µm).

**Figure S1** displays the dependence of the cells flow velocity, through the deformation microchannels, on various parameters such as the PEO concentration and molecular mass (in the suspending medium) and the inlet pressure. The linear velocity of Jurkat cells at the end of the deformation microchannels was determined by analyzing high-speed camera images. The linear flow velocity was evaluated for the following parameter sets: a) polymer concentrations of 0.1%, 0.5%, and 0.8%, b) pressures of 500, 1000, 1500, and 2000 mbar, and c) molecular masses of 1, 2 and 5 MDa. The tested conditions led to a considerable flow velocity variation between 5 cm/s and 200 cm/s. The flow velocity decreased monotonically and non-linearly with increasing PEO concentration and molecular mass, while it increased with higher inlet pressures. Under all these conditions, the cells were focused at the center of each focusing microchannel. Accordingly, the composition of the suspending medium and the applied pressure can be varied in a rather wide range without any major detrimental effect on the focusing efficiency. Moreover, we examined the dependence of cell deformation on the inlet pressure, the PEO molecular mass and the PEO concentration of the suspending medium (**Figures S2)**. For each parameter set in each diagram, 5000 cells were imaged during their passage through the deformation microchannels. Increasing pressure, polymer molecular weight, and concentration result in an upward shift of the deformability values.

### Blood and breast cancer cell lines phenotyping

To assess the capabilities of the proposed method for cell deformability analysis, we initially examined two subpopulations of blood cells, red blood cells (RBCs) in 20X diluted blood and peripheral blood mononuclear cells (PBMCs) (see Material and Methods for the isolation process). The samples were injected into the chip at a pressure of 1 bar and subsequently, characterized for their mechanical properties. The data for each cell type was acquired individually and is presented together in a scatter plot (**Figure 2a**). As shown in **Figure 2a**, although the two blood cell populations have similar sizes, they can be readily distinguished based on their differences in cell stiffness despite their comparable cell size. Specifically, red blood cells exhibited greater deformability, with an average value of 0.18, while PBMCs were found to be stiffer, with a mean deformability value of 0.04. Thus, the VDC approach successfully discriminated between RBCs and PBMCs.

**Figure 2.**
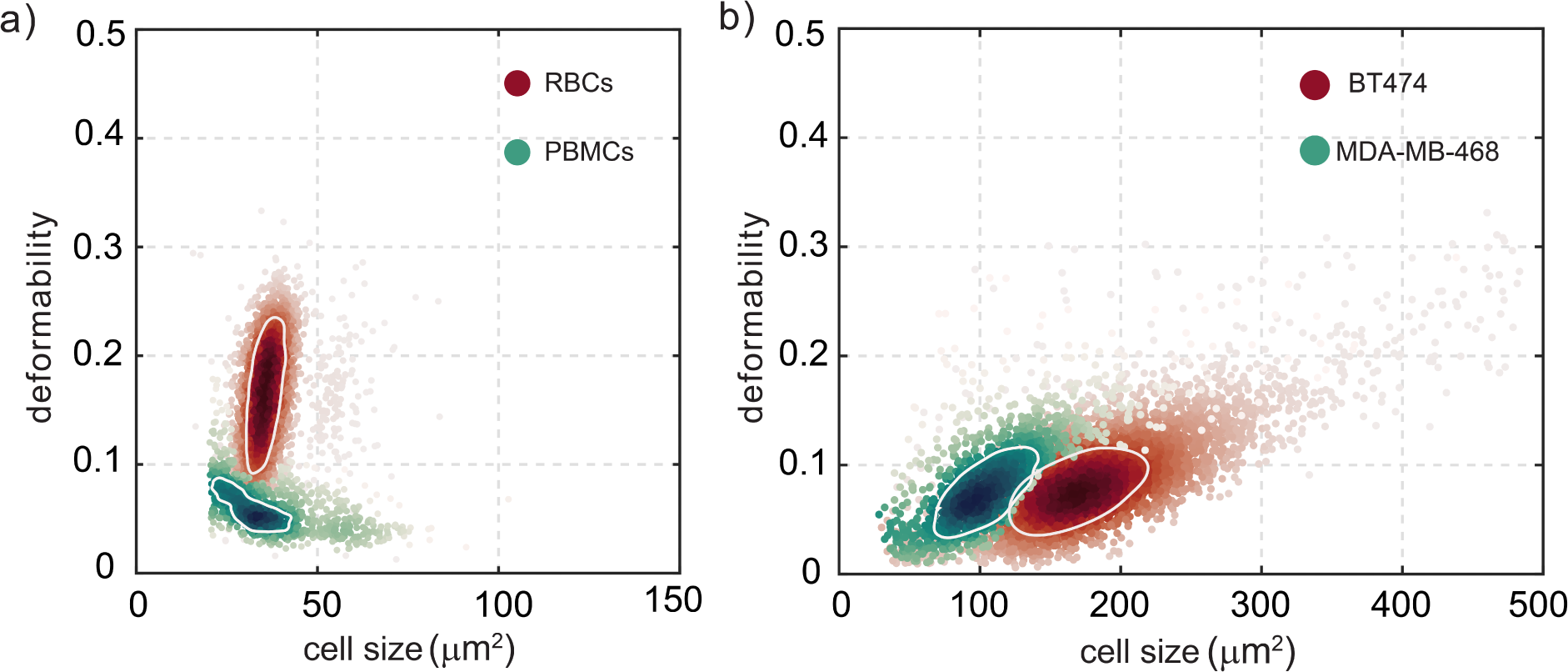
Mechanical phenotyping of blood and breast cancer cells. a) Density scatter plot of RBCs and PBMCs measured in separate experiments at the same pressure of 1 bar in a 15 µm x 15 µm channel but shown in one plot. The cells show differences in deformability while having similar sizes. b) Density scatter plot of breast cancer cell lines of BT474 and MDA-MB-468 measured in separate runs at the same pressure of 1 bar but shown in one plot. The cells have similar deformability levels while having different sizes. Both in (a) and (b) the white circles correspond to 50%-density contour plots.

Next, we investigated the mechanical phenotyping of two breast cancer cell lines, BT474 and MDA-MB468 as illustrated **in Figure 2b**. The cells were injected at a pressure of 1 bar and their passage through a deformation microchannel was imaged separately for each cell line in separate experiments. The comparison of deformability values revealed significant differences in cell size between the two cell lines, while their deformability remained similar.

### Effect of actin and tubulin drug on cell deformability

Herein we utilized VDC to investigate the effect of various pharmacological reagents on the mechanical properties of Jurkat cells. Previous research has demonstrated that the actin and microtubule networks significantly affect the mechanical properties of cells.^33, 34^ Hence, we employed cytoskeletal perturbing drugs, to alter these structures and assess their effect on cell deformation. Specifically, we used Latrunculin B (Lat B) and Cytochalasin D (Cyto D) to alter actin, and Nocodazole (Noco) to modify the microtubule network.^35, 36^ Actin is a structural protein of the cell cytoskeleton that forms the cell shape and morphology in its filamentous form (F-actin).^32^ Microtubules are long, rigid cylindrical biopolymers comprised of assembled tubulin dimers unveiled to resist contractile forces and interact with other cytoskeletal polymers to stabilize the cytoskeleton.^37^ Here, we utilized Cyto D and Lat B to prevent filament polymerization and Noco to stimulate the microtubule filament disassembly. For all experiments, an inlet pressure value of 500 mbar was used, and each set of experiments consisted of around 5000 cells per run, with three replicates for each set. Using VDC, we acquired cell size and deformation measurements of individual cells. Briefly, cells suspended in the viscoelastic medium were firstly introduced into the microfluidic device. As the cells travel through the focusing region, they aligned in single files in the parallel channels. Subsequently, the hydrodynamic forces in the constriction region deform the cells, and images are captured at the end of the deformation region. The 50% event density was used to generate a contour plot to facilitate the comparison. Lat B concentration was changed between 25 nM and 250 nM and compared with fixed and no-drug control measurements. **Figure 3a** displays the relationship between deformation and cell size for various Lat B drug concentrations, “no drug” (control), and fixed cells. The comparison of the contour plots reveals a considerable difference in deformability between fixed and viable cells. **Figure 3b** and **3c** illustrate bar graphs which represent the mean values of cell deformability and cell size as determined by the scatter plot. These measurements were performed with three replicates for each, and the chip was loaded with cells at four different drug concentrations. Quantitative analysis of the replicates reveals that the samples exposed to concentrations up to 125 nM did not show a significant shift in cell deformability. However, at a concentration of 250 nM a population shift toward elevated deformation values was observed as confirmed by analysis of variance (ANOVA), which shows a significant effect on the cell deformability (*p* = 0.0003). In the case of the cell size, no significant change was observed between the drug treatment conditions (*p* = 0.88-0.99). Similarly, the effect of Cyto D drug concentration on actin depolymerization was investigated using cell deformability and size measurements (**Figure 3d**). Results showed that at concentrations above 20 µM, the populations show an elevation in the deformation values (at 20 µM *p* <0.0001) (**Figure 3e-f**). However, no significant change in cell size was observed upon drug treatment. On the other hand, viable cells exposed to Noco showed higher deformability compared to fixed cells (**Figure 3g**). The exposure of control cells to Noco up to 10 µM did not show a significant increase or decrease of the mean cell deformation (*p* = 0.9903). However, a concentration of 100 µM Noco led to an upward shift of the population towards higher deformation values (*p* = 0.0033) while no significant change in cell size was observed upon drug treatment at all concentrations (*p* = 0.87-0.99). (**Figure 3h-i**).

**Figure 3.**
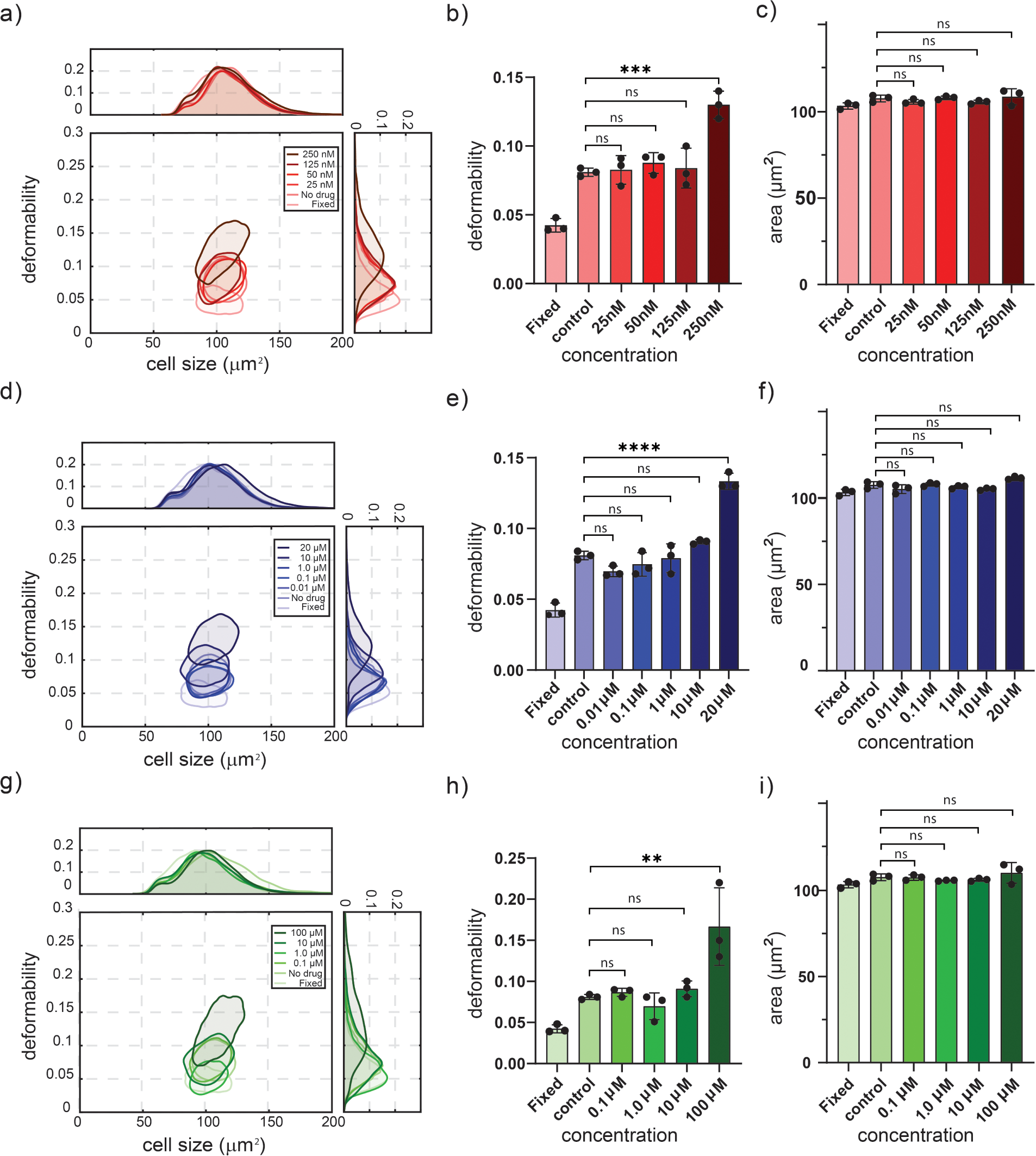
Effect of Lat B, Cyto D, and Noco drugs on cell deformation. a,d,g). Contour plots (50% density of the different subpopulations of deformability versus cell diameter for fixed Jurkat cells, cells incubated without (control) and with different concentrations of drugs. Measurements of deformability (b, e, h) and cell area (c, f, i) under different drug concentrations were carried out in multiple experimental replicates. Each dot on the bar graphs represents the mean value measured for 5000 cells and error bars represent standard deviation. For Lat B, Cyto D, and Noco, the concentrations that showed considerable deformation increase are 250 nM, 20 µM, and 100 µM, respectively. Statistical comparisons between these samples were performed using a one-way ANOVA; in alll cases the mean cell size difference between control and drug treated cells is not statistically significant. On the contrary deformability is significantly different from the control sample in the cases of 250 nM Lat B (*p* = 0.0003), 20 µM Cyto D (*p* <0.0001) and 100 µM Noco (*p* = 0.0033).

### Phenotyping of patient-derived glioma cells with cancer stem-cell properties vs. more differentiated glioma cells

Glioblastoma is the most common and most aggressive primary brain tumor in adults.^38^ Different cancer cell populations have been described in the context of glioblastoma, comprising glioma-initiating cells and more differentiated glioma cells.^39, 40^ Patient-derived glioma-initiating cell lines (GIC) have stem-like properties (tumor-initiating potential)^41^, are cultured in serum-free conditions and are more resistant to chemotherapy and irradiation compared to more long-term differentiated glioma cell lines (LTC) that are cultured under serum-containing conditions.^40, 42, 43^ The specific identification of glioma-initiating cells with cancer-stem cell properties remains a significant challenge because of their plasticity and due to a lack of selective markers. While some surface markers or combinations thereof have been associated with cancer-stem cell properties they are not exclusively specific to cancer stem cells.^44^ We used VDC to characterize the mechanical properties of GIC and LTC and compared them to non-malignant astrocytes. As shown in **Figure 4a (left panel)** the mean cell sizes of GIC cell lines, ZH-161, ZH-562, ZH-305, GS9 and GS5 are significantly different compared to those of LTC cell lines, LN-428, LN-18, LN-229 and T98G. In addition, all glioma cell lines have a different size compared to primary human astrocytes (control). On the other hand, while the deformability among all cell lines appeared to be quite heterogeneous (**Figure 4a, middle panel**), the size to deformability ratio presents a clear phenotype difference between the GIC (stem-like) and LTC (differentiated) cell lines (**Figure 4a, right panel**). In addition, the Kernel density plots of mean cell deformability against cell size revealed three different clusters resembling GIC, LTC and primary astrocytes (**Figure 4b left panel**). Owing to the information depth provided by images of phenotype parameters such as deformability and size (**Figure 4b, right panel**) VDC can differentiate distinct cancer cell populations in biopsy samples from tumours. Our results also show that cells from both types of tumours had significantly higher mean cell size and size to deformability ratio than their healthy counterparts (**Figure 4c)**.

**Figure 4.**
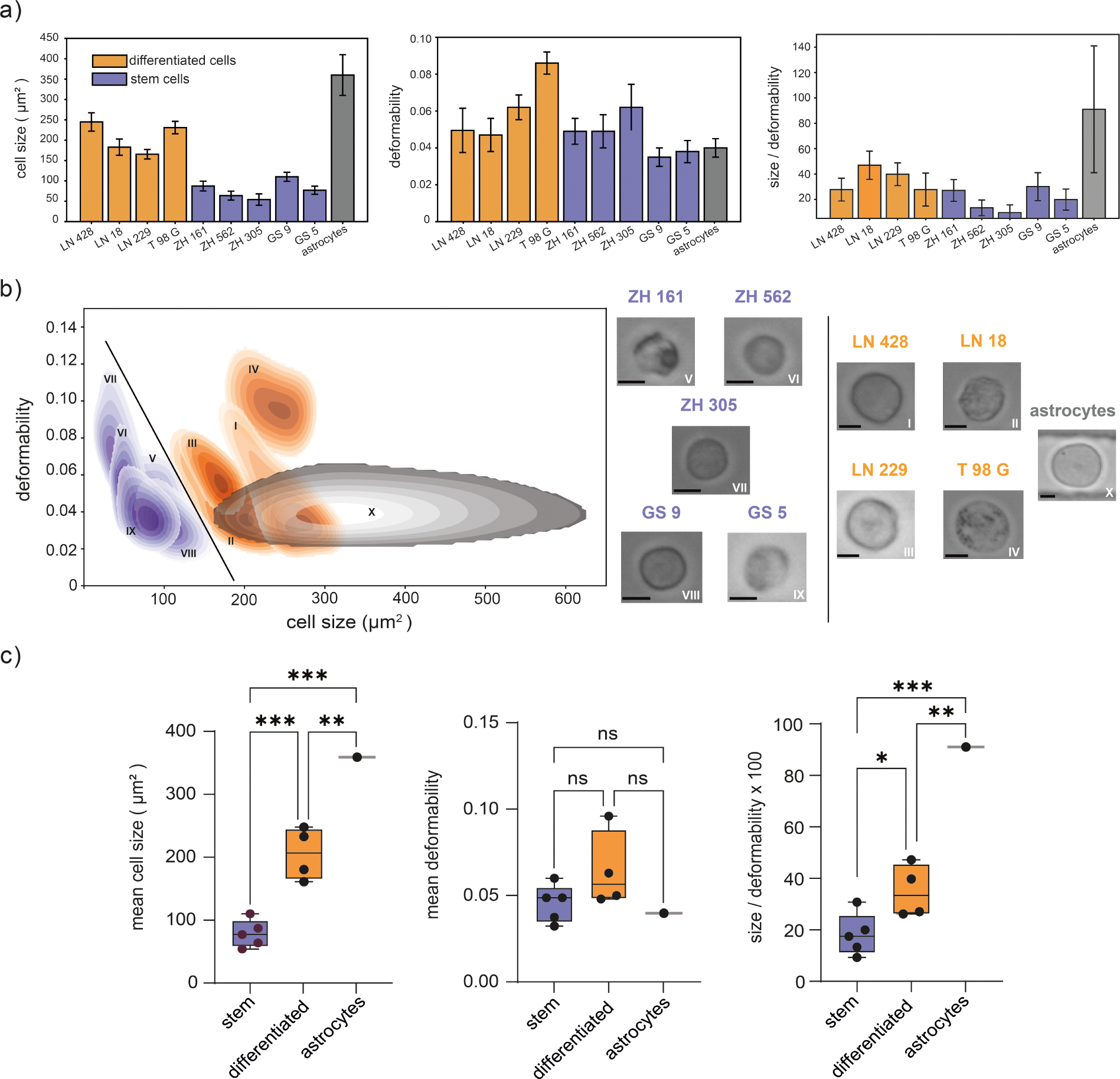
Investigation of glioblastoma stem and differentiated cell signatures using viscoelastic deformability cytometry. a) Bar charts depicting the mean cell sizes and deformabilities of differentiated and stem-type glioma cancer cells. Error bars represent standard deviations derived from three independent experiments. b) All the tested types of glioma cells can be identified on the basis of joint Kernel density estimate plots of deformability vs cell size (left part). Two clusters of cells (divided by the solid line) showing that the two types of glioma cell lines GIC (purple) and LTC (orange) can be identified on the basis of the density plots, while the control cell line (astrocytes: dark gray) shows greater heterogeneity of cell size compared with the solid tumor derived glioma cells. Brightfield images (right panel) of both differentiated- and stem-type glioma cells where features such as size and deformability can be extracted for multi-dimensional analysis (scale bar: 10 µm). c) Box graphs of size, deformability, and their ratio between two different types of glioma cells (GIC: purple, LTC: orange) and a control cell line (dark gray). The horizontal line represents the mean value. Box plot whiskers range from 5th to 95th percentiles and the length of the whiskers is restricted to a maximum of 1.5 times the interquartile range. Statistical comparisons between these samples were performed using a one-way ANOVA; Mean cell size difference of stem vs. differentiated (*p* = 0.0009), stem vs astrocytes (*p* = 0.0001) and differentiated vs. astrocytes (*p* = 0.0059); The mean deformability value among the above samples is not significant (ns); Mean cell size to deformability ratio of stem vs. differentiated cells (*p* = 0.0476), stem cells vs astrocytes (*p* = 0.0003) and differentiated cells vs. astrocytes (*p* = 0.0018) show that these three different categories are statistically significant.

### Chronic lymphocytic leukemia diagnosis using VDC

As mentioned previously mechanical phenotyping through the measurement of cell deformability represents a potentially powerful diagnostic tool. Since cells undergo physical and biological changes during cancer progression, a complete understanding of such changes offers much in developing new diagnosis and treatment methods. Leukemia is a malignant disease of the hematopoietic system, including bone marrow, with survival rates dropping from 90% for good-risk diseases to 20% for poor-risk diseases.^45^ A contemporary leukemia diagnosis is mostly based on a combination of conventional microscopy, flow cytometric detection of fluorescent cell surface markers, and genetic analysis.^46^ Such diagnostic workflows are complex, costly, and time-consuming; thus, label-free, cost-effective, and rapid single-cell analysis techniques are needed to enable the diagnosis of hematological malignancies in a robust manner. Previously deformability cytometry has been used as a label-free marker to diagnose malignant pleural effusions, including discriminating leukemias from inflammatory processes.^23^ However, no study so far has shown the potential of mechanical phenotyping to diagnose leukemia in a clinical setting.

To tackle this issue, we employed VDC for high throughput extraction of malignant cells deformability from both healthy and leukemia-affected human blood samples. In particular, we focused on classifying Chronic Lymphocytic Leukemia (CLL) cells by mechanically characterizing a large number of suspended cells obtained from patients with lymphoma (a type of malignant B) and compared them with peripheral blood mononuclear cells (PBMCs) and B-cells from healthy donors. **Figure 5a** displays a scatter plot that encompasses these scenarios, revealing that healthy and malignant (CLL) cells can be differentiated by their size and deformability, with two distinct regions corresponding to the two cell types. Further analysis carried out on cells obtained from a cohort of healthy donors and leukemia patients demonstrates considerable differences in both cell size and deformability. Specifically, 10 PBMS samples and 10 B-cell samples from healthy donors, as well as 10 CLL samples, were analyzed using VDC. **Figure 5b (left panel)** shows that CLL cells have higher deformability values compared to healthy PBMC and B cells, indicating a potential diagnostic application for the VDC technique in distinguishing between healthy and malignant cells. Statistical t-test analysis showed that the mean cell deformability of CLL is 0.09, significantly higher (*p* < 0.0001) that of the PBMCs and B cells which yield a mean value of 0.05 (**Figure 5c**). In addition, the analysis of all samples revealed that CLL cells had also significantly higher mean cell area compared to healthy PBMCs (*p* < 0.0298) and B cells (*p* < 0.0001) (**Figure 5c**). Based on the results, we conclude that VDC can precisely determine normal and abnormal B cells.

**Figure 5.**
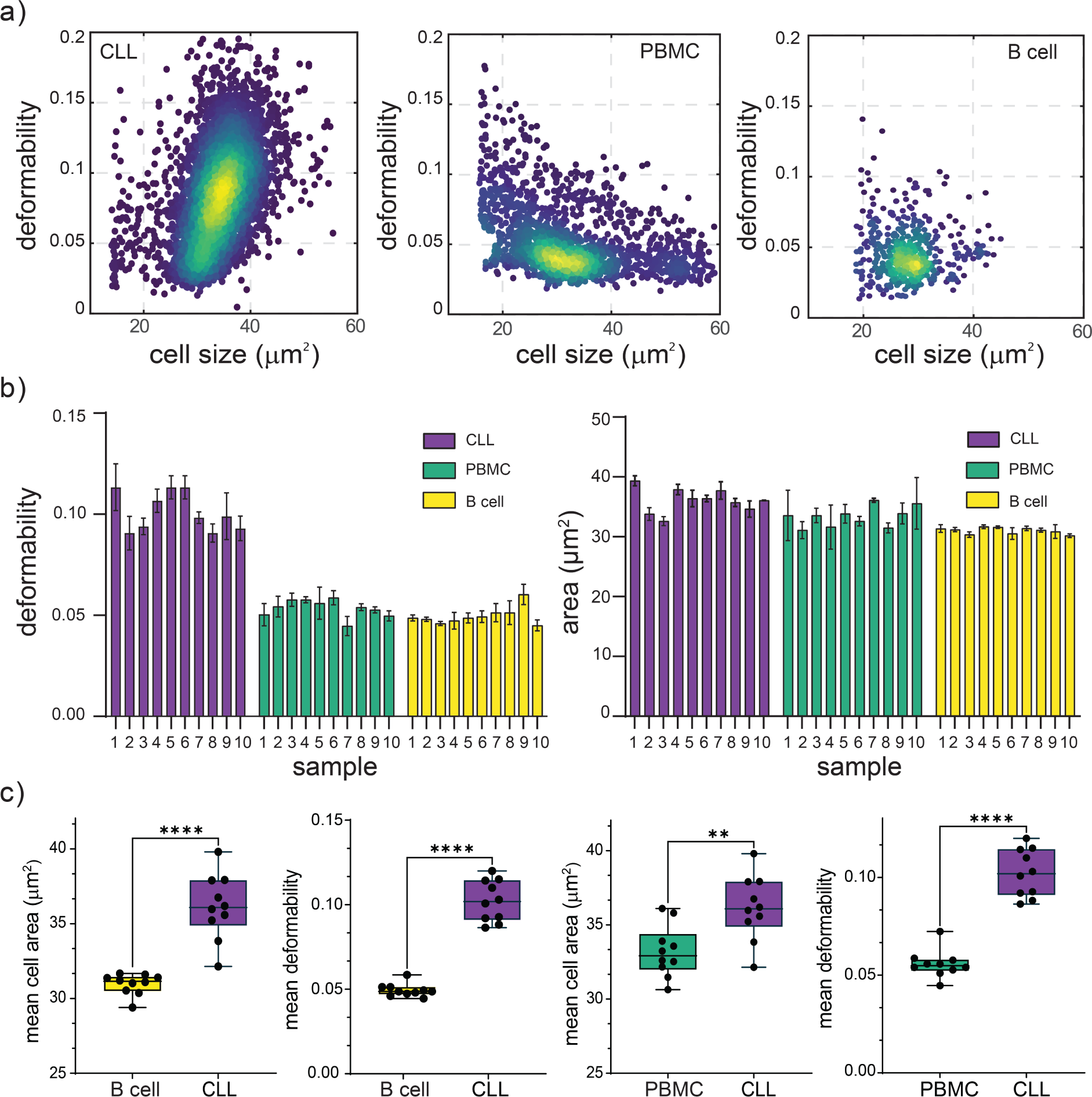
Investigation of CLL diagnostic signatures using viscoelastic deformability cytometry. a) Deformability versus cell size density scatter plots for samples containing CLL, healthy PBMCs, and healthy B cells. b) The comparison of deformability and size was measured for 10 healthy PBMC, 10 healthy B cell, and 10 CLL samples. For each sample, at least three replicates were measured, each one having 5000 cells and the mean values were considered. c) Box graphs of mean cell deformability and size of control (green) and lekeumia samples (purple). The horizontal line represents the means of these phenotype parameters. Box plot whiskers range from 5th to 95th percentiles and the length of the whiskers is restricted to a maximum of 1.5 times the interquartile range. Statistical comparisons between B cells, PBMCs and CLLs were performed using a two-tailed Mann-Whitney U test; B cells vs CLLs mean have a significant difference in cell size (*p* < 0.0001), and deformability (*p* < 0.0001); PBMCs vs CLLs have a significant mean cell size difference (*p* = 0.0089) and deformability (*p* < 0.0001). CLL samples show significantly higher mean deformability and mean size values than the control (healthy) B-cell and PBMC samples.

## Conclusions

To conclude, we have introduced an ultra-high throughput microfluidic platform that enables the measurement of cells’ mechanical behavior under stress. Owing to the viscoelasticity of the fluid, cells can be focused and deformed in the same device, eliminating the need for any sheath fluid. Treatment of cells with Lat B, Cyto D, and Noco reagents, leads to depolymerization of the actin and tubulin filaments, causing changes in cell deformability. Furthermore, we utilized VDC to mechanically phenotype cells and identify diseased populations in clinical samples. Specifically, we showed that VDC can differentiate cancer cells with stem cell properties from more differentiated cancer cells in the context of glioma type cells and enables the diagnosis of lymphoma based on single-cell bright-field imaging of PBMCs. More specifically the results revealed elevated deformability values for CLL in comparison to healthy PBMCs and B cells.

The deformability platform presented here can be applied directly for the identification of diagnostic signatures for various diseases using blood or other body fluid samples or tisse samples. We expect that our approach, will circumvent the challenges associated with defining molecular diagnostic markers and rather allow diagnosis of hematological or solid tumor malignancies owing to its rapid, high throughput, automated capabilities. Such an advance can allow, the ultra-high throughput and sensitive diagnosis of several diseases, as well as rare event detection and whole blood cell classification. For example, altered cellular deformability is a component of multiple clinically significant red cell abnormalities, such as Sickle Cell Disease or Thalassemia^47-49^. In this regard, it is noted that osmotic gradient ektacytometry is a valuable screening test for hereditary hemolytic anemias.^50^ However, ektacytometry is technically demanding and established in only a few hematological laboratories worldwide. Deformability measurements using the developed microfluidic platform require only minute amounts of whole blood, are cost-effective to perform and high-throughput in nature. Accordingly, they can be easily and widely implemented in hospitals and clinics, rather than existing in centralized testing facilities, as is currently the norm for flow cytometry instrumentation. The automated detection of PBMC cells will allow accurate and precise diagnosis of lymphoma and leukaemia and will facilitate the diagnostic exclusion of clinically similar inflammatory diseases and infections. Furthermore, compared to manual microscopy, the higher sensitivity and specificity of image cytometry will be beneficial in the assessment of the dynamics of lymphoma or leukemia during therapy and monitoring the recurrence of the disease for patients in remission or after chemotherapy.

The presented image-based viscoelastic platform when combined with high-speed imaging is able to measure cell deformation at rates up to 100,000 cells per second. Notably, the platform was able to detect malignant cancer leukemia cells from a cohort of cancer patients and discriminate them from a cohort of healthy samples. This is of particular importance for clinical studies, where millions of cells must often be analyzed in order to detect rare cancer cells. Additionally, it should be noted that currently there exists no competitor technology able to image and process cells at throughputs in excess of 1000 cells per second. Put simply, the integration of image-based deformability cytometry and viscoelastic microfluidics represents a paradigm shift in flow cytometry technology for the diagnosis of hematological diseases and solid tumors. In principle, clinical application of this technology is straightforward, and thus we expect that clinicians will be able to directly use deformability signatures in their clinical workflows. Furthermore, although we focused on lymphoma diagnosis and different types of glioma cell identification, the deformability cytometer can be used for the diagnosis of a variety of diseases. VDC has the potential to open new opportunities in the assessment of blood disorders, since diagnostic information is generated in a rapid and automated fashion. Using such an approach, the challenges associated with defining molecular diagnostic markers may be circumvented, which in turn will resurge deformability-based diagnosis of diseases in a sensitive, affordable, and high throughput manner. Put simply, since the physical properties of cells (such as cell deformability and cell/nuclear size) are promising label-free biomarkers for cancer diagnosis and prognosis, the development of tools for their measurement provides a valuable advance in personalized medicine.

## Materials and Methods

### Microfluidic device fabrication

The fabrication process of a microfluidic device has been schematically illustrated in **Figure S3**. The microfluidic device has two sections with two different channel heights. The deformation section comprised ten parallel rectangular channels of 300 μm length, 15 μm width and 15 μm height, and the focusing section comprised ten parallel channels of 3 cm length, 50 μm height and 50 μm width. We utilized an SU8 negative photoresist for creating the master mold by means of standard photolithographic techniques. To account for the different heights of the two device sections, which were 50 μm and 15 μm, the master mold was fabricated in two steps by depositing and patterning two-layer portions with different heights onto a silicon wafer.

For fabricating the master mold, two masks were designed using AutoCAD 2019 (Autodesk, San Rafael, CA, USA) and were laser-printed on a 5-inch Cr/fused silica transparency mask (Micro Lithography Services Ltd, Chelmsford, UK). The first mask served to fabricate the deformation section, and the second mask served to fabricate the focusing section. The first layer portion was fabricated as follows: SU-8 2010 photoresist (PR) was spin-coated (acceleration: 500 rpm/s, speed: 1500 rpm, time: 30 seconds) on the silicon wafer (4-inch diameter) and prebaked at 65 °C for 3 minutes and 95 °C for 9 minutes. Next, the photoresist was exposed to UV light at 140 mJ/cm² intensity, using the first mask, and post-baked at 65 °C for 2 minutes and 95 °C for 4 minutes. The photoresist was developed for 3 minutes and hard-baked at 150 °C for 10 minutes. This resulted in the first layer portion, having a height of 15 μm (**Figure S3a)**. Next, the second mask was aligned with the first layer portion using mask aligners. The second layer portion was fabricated as follows: SU-8 2050 photoresist (PR) was spin-coated (acceleration: 500 rpm/s, speed: 3000 rpm, time:30 seconds) on the previously prepared silicon wafer and prebaked at 65 °C for 3 minutes and 95 °C for 9 minutes. Subsequently, the photoresist was exposed to UV light at 150 mJ/cm² intensity, using the second mask, and post-baked at 65 °C for 2 minutes and 95 °C for 4 minutes. The photoresist was developed for 5 minutes and hard-baked at 150 °C for 10 minutes. In this manner, the second layer portion, having height h_1_ = 50 μm, was created side-by-side with the first layer portion (**Figure S3b)**.

For producing the microfluidic device, a 10:1 mixture of polydimethylsiloxane (PDMS) monomer and curing agent (Sylgard 184, Dow Corning, Midland, MI, USA) was poured over the master mold and polymerized at 70 °C for 4 h (**Figure S3c)**. Inlet and outlet ports were punched using a hole-puncher (SYNEO, West Palm Beach, FL, USA) **(Figure S3d)**. The structured PDMS substrate was then bonded to a 1 mm glass substrate after treating both surfaces in an oxygen plasma (EMITECH K1000X, Quorum Technologies, Lewes, UK) for 60 seconds (**Figure S3e)**. The finished device is illustrated in **Figure S3f**.

### Cell culture and sample preparation

Jurkat (Sigma-Aldrich, Buchs, Switzerland), BT474 (ATCC, Manassas, Virginia, USA), and MDA-MB468 (ATCC, Manassas, Virginia, USA) cell lines were cultured in polystyrene flasks using Gibco DMEM high glucose medium (ThermoFisher Scientific Inc., Waltham, Massachusetts, USA) and Gibco RPMI 1640 medium (ThermoFisher Scientific Inc., Waltham, Massachusetts, USA) mixed with 10% (v/v) FBS (Life Technologies, Zug, Switzerland) and 1% (v/v) penicillin– streptomycin (Life Technologies, Zug, Switzerland). LTC (LN-428, LN-18, LN-229, T98G) and GIC (ZH-161, ZH-562, ZH-305, GS9, GS5) glioma cell lines have been described before and were cultured as reported before.^51^ Flasks were stored in an incubator (New Brunswick Galaxy 170S, Eppendorf, Basel, Switzerland) at 37 °C, 5% CO_2_, and 95% humidity, and the medium was refreshed every 2 days. For all experiments, cellular concentrations were maintained at 2 million cells per mL. For each experiment, cells were centrifuged (120 rpm for 5 minutes at 24 °C), washed in PBS, filtered through a strainer with a pore size of 40 μm (Corning Inc., New York, NY, USA), centrifuged, and re-suspended in the PBS. Treatments to modify cell stiffness via cell fixation and administration of Latrunculin B (Lat B) and Cytochalasin D (Cyto D), and Nocodazole (Noco) were performed as follows: Cyto D, Lat B, and Noco were dissolved in dimethyl sulfoxide (DMSO) according to standard protocols.^18, 32, 33^ The following concentrations were used to investigate the dose-response: 0.01 μM, 0.1 μM, 1 μM and 10 μM of Cyto D; 0.1 μM, 1 μM, 10 μM, and 100 μM of Noco; and 25 nM, 50 nM, 125 nM, and 250 nM of Lat B. The DMSO was dissolved at a low concentration of 0.1% in all experiments to minimize the DMSO effect on cell deformation. Fixation was performed by the addition of a 4% paraformaldehyde solution for 10 minutes at room temperature. To prevent the formation of cell aggregates, cells were subsequently washed with PBS supplemented with 10% FBS. For making viscoelastic solution, polyethylene oxide polymers (M_w_: 1 MDa, M_w_: 2 MDa, and M_w_: 5 MDa, Sigma-Aldrich, Buchs, Switzerland) were fully dissolved in PBS to a concentration of 1% (w/v). The prepared solution was then aged for one week at 4°C to reach steady-state viscosity. Finally, the PEO solution was added to the cell medium at concentrations of 0.1%, 0.5%, and 0.8% for the deformability experiments.

### PBMC isolation

Mononuclear cells from healthy human whole blood and CLL patients were isolated by density centrifugation using SepMate-50 tubes (STEMCELL Technologies, Vancouver, Canada). In short, a density gradient medium (1.077 g/mL) was added to the tubes through the insert, and blood samples, previously diluted 1:2 with phosphate-buffered saline (PBS) and supplemented with 2% fetal bovine serum (FBS), were added. Density centrifugation was performed at 1200×g for 10 minutes at room temperature. Then, the top fraction (containing the enriched mononuclear cells) was removed, washed twice with PBS at 300×g for 8 minutes, and finally re-suspended in PBS.

### Deformability set up analysis

A pressure pump (Flow EZ, Fluigent, Le Kremlin-Bicêtre, France) was utilized to motivate the cells into the microfluidic device. High-speed brightfield imaging of cells was performed using a CB019MG-LX-X8G3 high-speed camera (XIMEA, Munster, Germany) mounted on a Ti-E microscope (Nikon, Zurich, Switzerland) using a 20×, 0.45 NA S plan Fluor objective (Nikon, Zurich, Switzerland), and a high-power collimated LED (UHP-T-560-DI-DF, Prizmatix, Holon, Israel) source. For a reduced region of interest (ROI) (1024 x 80 pixels), images were recorded acquisition at 10,000 f.p.s and in order to record blur-free images, the camera was operated at an exposure time of 1 or 2 μs. Images captured by the high-speed CMOS camera were transferred to a Linux multi-core PC using a camera-link frame grabber card.

### Deformability image analysis

All image processing steps are carried out using a Linux multi-core PC with a custom-written software. The image processing algorithm was implemented in a Python environment and in the real-time mode the algorithm was capable of performing image acquisition, image analysis and data storage for several hundred cells per second in real time using the OpenCV computer vision library (http://opencv.org). The image processing consists of the following steps: (i) background subtraction, (ii) threshold filtering, (iii) contour finding, and (iv) contour processing for the estimation of cell size, deformation, and brightness. More specifically for a selected ROI, the video is cropped, and the background image is calculated by averaging the intensities of the captured images. First, a median filter is used to reduce the noise. A binary filter with a specified threshold was used to detect cell contours. Then, for each cell, the cell size was determined, being the area A of the cell in the microscope image. In addition, the length l of the cell perimeter was determined. A dimensionless deformation parameter (called “deformability”) was calculated, using the formula

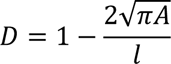

A two-dimensional scatter plot was created, using the size and deformation parameter of the cells, and the density of the points in the scatter plot was evaluated to determine a single contour line at 50% density. In addition, other physical parameters can be calculated from the images in real time, namely brightness and aspect ratio. Brightness is defined as the mean of all pixel intensities inside the cell contour and aspect ratio is the ratio of the large to the small axis of a cell. The software also calculates the area ratio defined as the convex hull area to the measured cell area and is used to indicate the presence of a cell (**Figure S4**).

The software architecture has a multi-thread structure whilst communication between different threads is achieved through queues. The threads would receive the data packets and calculate all parameters in real-time. When a cell passes through the constriction channel, the camera measures and streams the video frame to the threads of the software. In practice, the first thread is responsible for acquiring single frames from the camera and fetching them into a circular buffer. Then, the second thread is responsible for performing the image processing tasks that include background subtraction, thresholding to create a binary image, cell contour detection and calculation of the cellular attributes such as cell area, size and deformability. The processed data packets including scatter plot and images of the detected cell contours would then be sent to the GUI for display which are updated continuously using the third thread. The GUI would handle user inputs and configuration, such as selecting the region of interest. A scatter plot of cell size vs deformability is depicted in real time in the GUI (**Figure S5**) and also stored together with results from image processing such as brightness, aspect ratio and an image of the cell on the disk of the computer for later analysis. The processing time of a single frame for extracting all these parameters is between 0.5 and 1 ms

### Statistical Analysis

All statistical analyses were performed with GraphPad Prism 9.4 (GraphPad Software Inc., La Jolla, CA, USA). In all graphs the significance level was defined as p < 0.05 and p values are represented by *p < 0.033, ** p < 0.0021, *** p < 0.0002 and **** p < 0.0001. Statistical significance of overall differences of cell size and deformability data derived from the drug-treated and glioma cell experiments were analyzed by one-way analysis of variance (ANOVA) followed by Tukey’s multiple comparison test. CLL, B- and PBMC cell deformability and cell size data were analyzed by a two-tailed Mann–Whitney U test. This t-test was used to assess unpaired samples of healthy versus blood cancer patients. The p values reported from each experiment come from comparison of the given sample to the control condition obtained the above t-tests for multiple comparison.

### Numerical simulations

To evaluate the distribution of hemodynamic shear stress on the cell surface according to the viscoelasticity of the considered fluid, computational fluid dynamics (CFD) simulations were performed (**Figure S6)**. A representative geometrical model of a cell circulating in the deformation channel (width and height 15 µm, cell diameter 10 µm) was created for the purpose. Following a prior mesh independence study, the geometry was discretized into ∼1.05 million polyhedral and prismatic (five layers) elements. Steady-state, laminar simulations of the four PEO concentration conditions were performed in Ansys Fluent (v. 2020, Ansys Inc., Canonsburg, PA, USA). The boundary conditions were derived from the experimental setup of the considered scenario. Specifically, the mean velocity imposed as flat profile at the inlet section was derived from the relative flow-rate measurements at the inlet of the microfluidic chip, following the assumption of quantity conservation. At the outlet section, a relative zero-pressure was assumed. The no-slip condition was imposed on the rigid walls of the channel and the cell. Preliminary comparisons of the model of fluid were carried out comparing Newtonian and non-Newtonian Carreau models, assuming a constant density of 1080 kg/m^3^ (**Figure S6a and S6b**). For each PEO concentration, rheological data were extracted and introduced in Prism (v. 9, GraphPad, San Diego, CA, USA) to derive the relative parameters of the Carreau model. The other simulation settings are summarized in Table 1.

**Table 1.** Solver settings of the computational simulations.

| Type | Pressure-based Ansys Fluent |
| --- | --- |
| *Pressure velocity coupling method* | coupled |
| *Spatial discretization scheme* |  |
| gradient | least squares cell based |
| pressure | second order |
| momentum | second order upwind |
| *Steady state formulation* | second order implicit |
| ***Relaxation factors*** |  |
| *Flow Courant number* | 50 |
| *Explicit relaxation factors* |  |
| momentum | 0.35 |
| pressure | 0.35 |
| ***Convergence criterion residuals*** |  |
| continuity | $5 \times 10^{-5}$ |
| velocity | $5 \times 10^{-5}$ |
| max iterations | 1000 |

