## Supplementary Information for "Viscoelastic Deformability Cytometry: Ultra-high Throughput Platform for Mechanical Phenotyping of Cells in Liquid and Solid Biopsies"

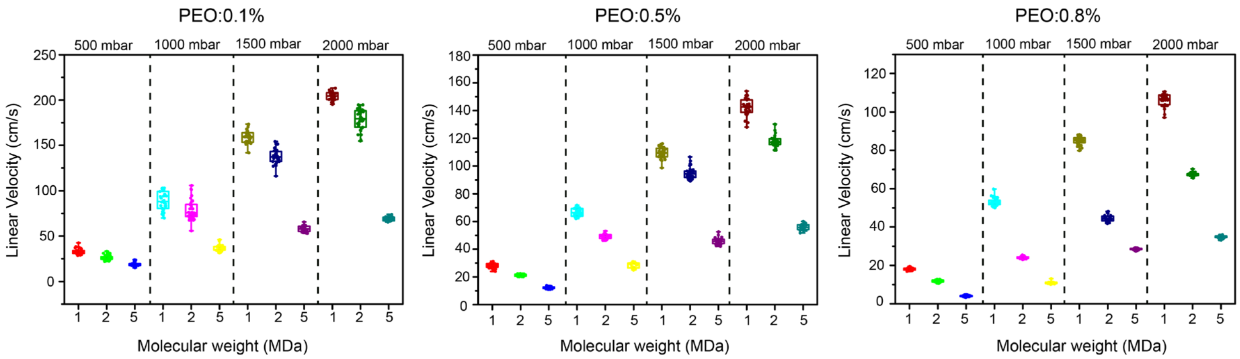


**Figure S1.** Plots of linear velocities of cells when suspended in three different PEO concentrations (1%, 0.5%, 0.8%) against PEO molecular weights of 1, 2, and 5 MDa at inlet pressure values of 500, 1000, 1500, and 2000 mbar. For each analysis, the velocity of 30 cells were measured at the end of the deformation region.

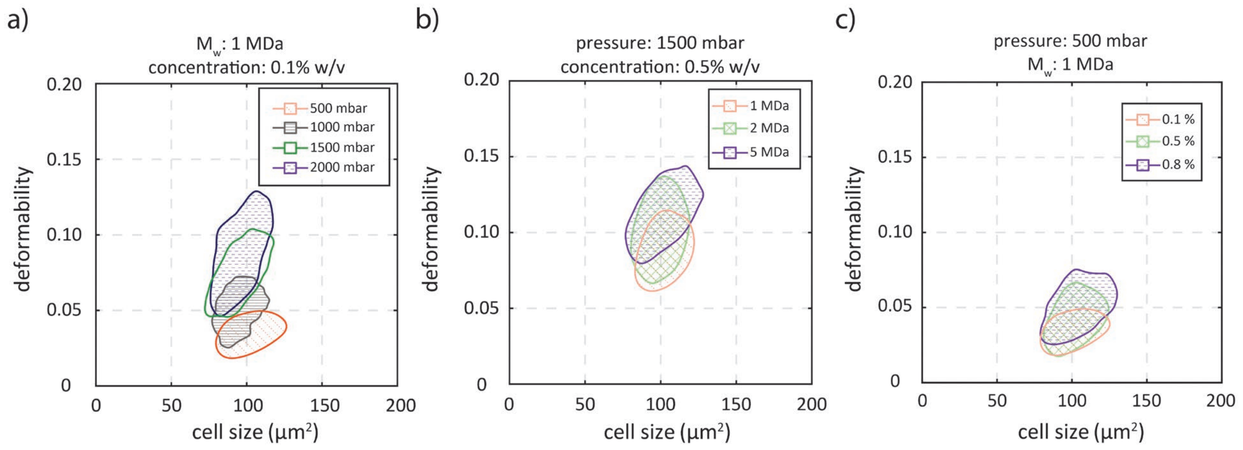


**Figure S2. Scatter plots of deformability vs cell size under the influence of pressure, PEO molecular and concentration**. The effect of inlet pressure (a) PEO molecular weight (b) and PEO concentration (c) on cell deformation.

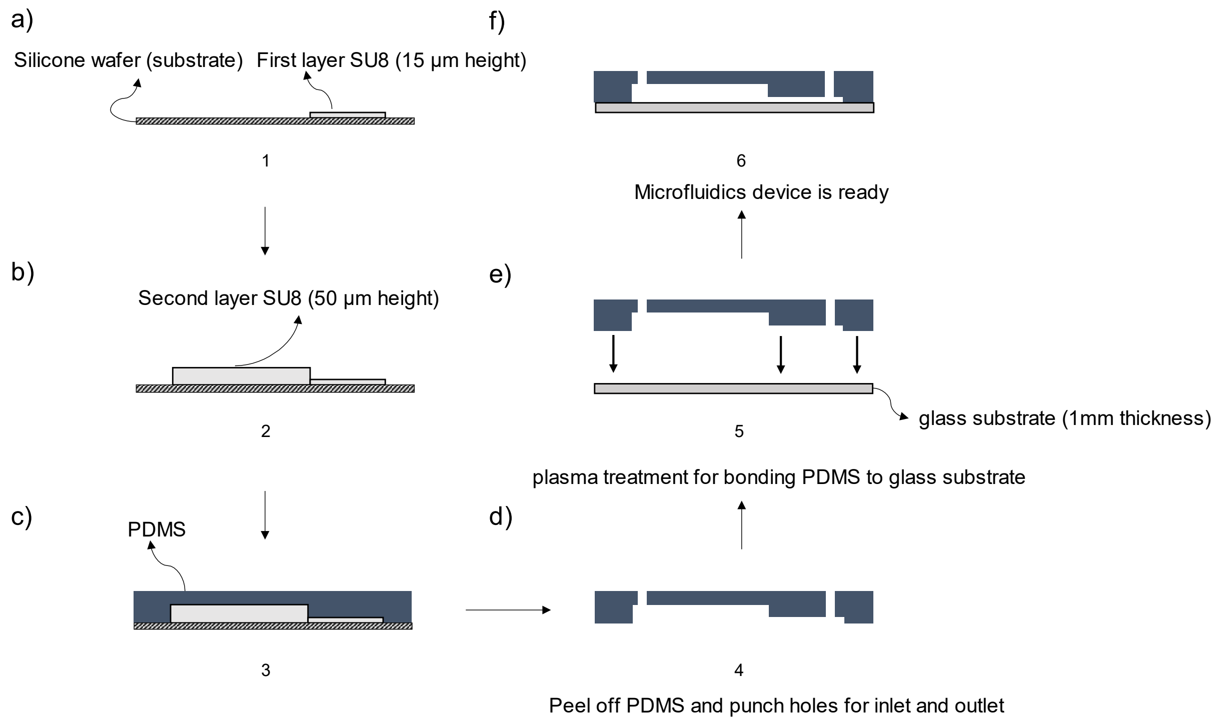


**Figure S3.** Fabrication process of the two-layer PDMS microfluidic device. a) the first SU8 layer with a height of 15 µm is fabricated on the silicon wafer. b) The second SU8 layer, having a 50-µm height is aligned next to the first layer and subsequently fabricated. c) PDMS is casted on the multilayer mold and subsequently cured. d) PDMS is peeled off and inlet and outlet ports are created. e) The structured PDMS is plasma-bonded on a glass slide. f) The final state of the completed microfluidic device.

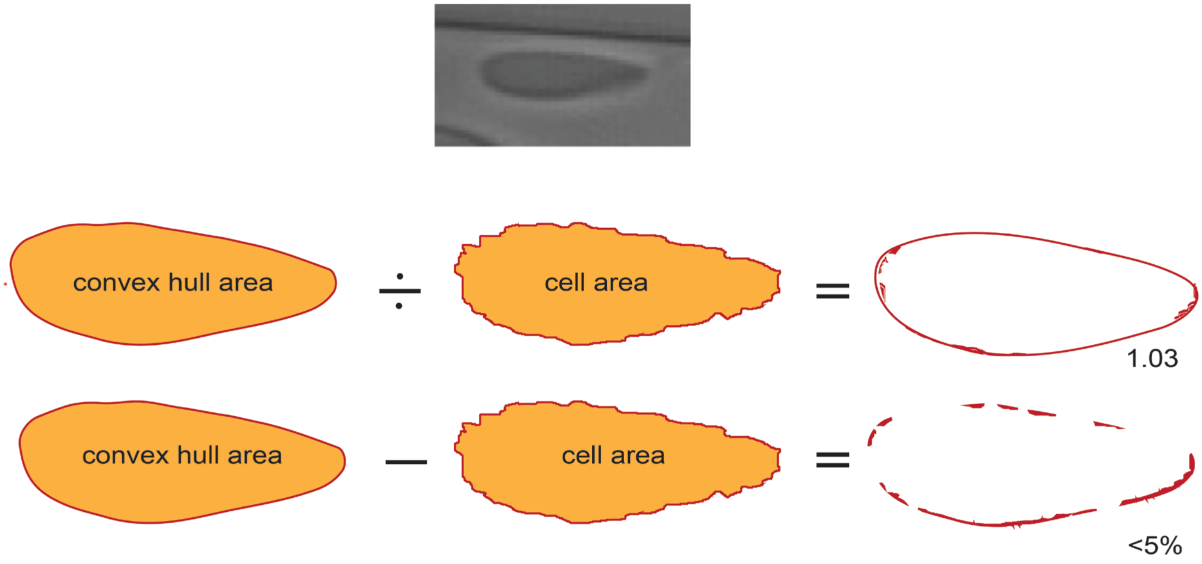


**Figure S4.** Graphical representation of the area ratio and difference. Cells with an area ratio below 1.05 or an area difference less than 5%, have a convex contour, whereas cells with area ratio above 1.05 or an area difference more than 5% are filtered out. Exemplary image of a cell with an area ratio of 1.03.

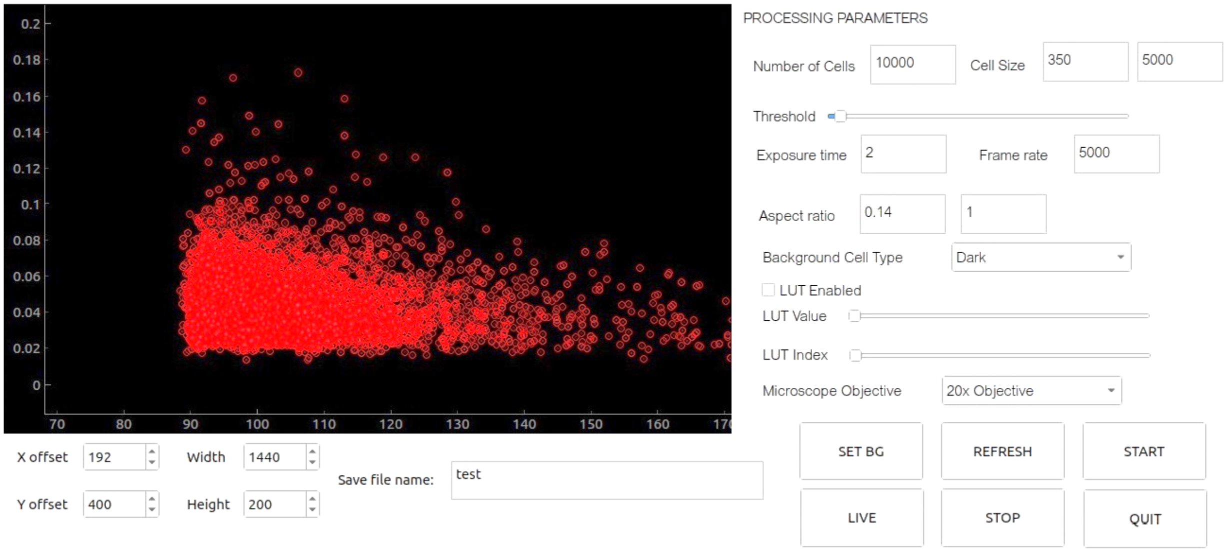


**Figure S5. Screen capture of the GUI of the video processing software for real-time measurements of various metrics**. Inputs parameters such as the number of cells to be analyzed, the cell size, the threshold as criterion of image binarization, exposure time and frame rate of the camera, the cell aspect ratio and the image background type can be user defined. Results are displayed in a 2D scatterplot, showing the deformation parameter and the cross-sectional area of single cells. Each dot represents the result of a single cell measured out of a total thousand cells. For each cell detected additional parameters such as the cell size and the mean brightness are exported for further analysis.

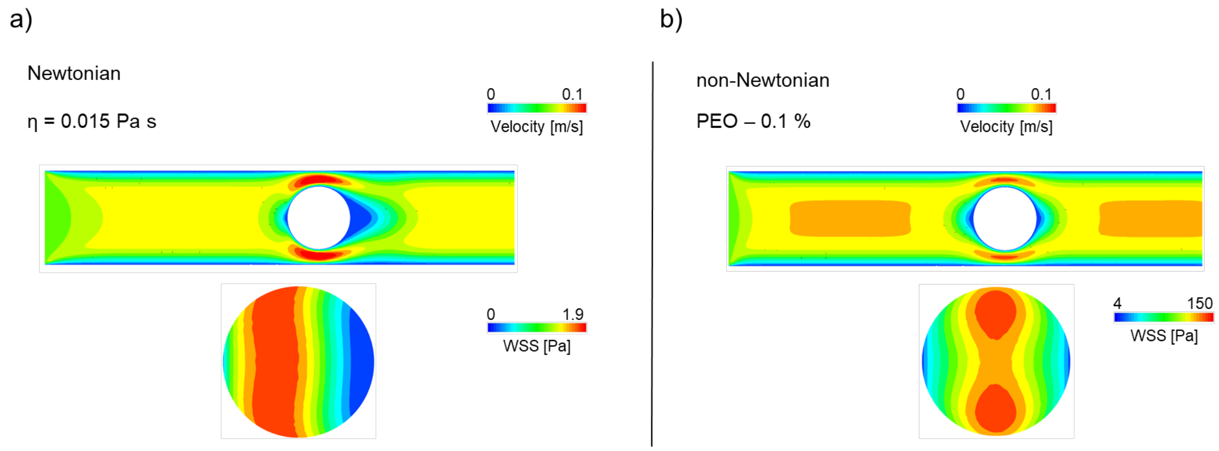


**Figure S6. Analysis of the impact of the fluid properties on the wall shear stress (WSS), under the no slip boundary conditions**. In a 15 µm channel, a cell with a diameter of 14 µm was positioned at the center. Simulations were conducted using Newtonian fluid (a) and 0.1% PEO 1MDa (b) viscoelastic fluid to determine the distribution of wall shear stress around the cell. A fixed flow rate of 3 µl/s was used for both conditions. WSS values and their distribution patterns are different for Newtonian and viscoelastic fluids.
